# A dataset of speed-resolved blood perfusion and oxygen saturation in human skin response to thermal stimulation

**DOI:** 10.1101/2020.03.04.976456

**Authors:** Shuyong Jia, Xiaojing Song, Weibo Zhang, Guangjun Wang

**Affiliations:** Institute of Acupuncture and Moxibustion, China Academy of Chinese Medical Sciences, Beijing, China

**Author notes:** corresponding author(s): Guangjun Wang.

## Abstract

The functional status of vessels can be determined by assessing blood perfusion. By integrating diffuse reflectance spectroscopy (DRS) and laser Doppler Flowmetry (LDF), the speed-resolved blood perfusion and blood oxygen saturation can be measured simultaneously by Enhanced Perfusion and Oxygen Saturation (EPOS). The dataset presented in this descriptor contains EPOS data recorded from a forearm point exposed to different levels of thermal stimulation, the classical LDF at control points, the R-R time series and data regarding characteristics of subjects. All data were recorded from 60 recruited healthy subjects. Half of the subjects received different levels of thermal stimulation, and half of them were blank controls. We believe that this dataset will lead to the development of local blood perfusion methods that can be used to index vessel function assessments. This publicly available dataset will be beneficial to the microcirculation evaluation.

## Background & Summary

There is increasing evidence that microcirculation can be used to evaluate vascular disorders at the systemic level[1, 2], and there is a close relationship between cardiac function and vessel function[3, 4]. In general, the functional status of vessels can be assessed by laser Doppler flowmetry (LDF). However, differentiating between different vascular compartments using the classical LDF approach is difficult. Recently, a multiparameter model based on the Monte Carlo algorithm has provided the possibility of further distinguishing the different velocity components in microcirculation perfusion[5–7]. This new method may provide further insight into evaluating vascular dysfunction at the systemic level[8, 9].

According to previous studies, blood perfusion signals recorded by LDF can be separated into different frequency bands in the frequency domain[10–13]; these frequency bands might reflect different physiological rhythms[14]. From another perspective, microcirculation perfusion can also be distinguished by the speed distribution, and the speed of red blood cells (RBCs) is closely related to the total cross-sectional area of different types of blood vessels. Therefore, different velocity components have potential value for distinguishing different types of blood vessels, thereby exploring the regulatory mechanisms of local microcirculation. By applying computational models, the Enhanced Perfusion and Oxygen Saturation (EPOS) system enables absolute measures of speed resolved perfusion. Dividing the flow in different speeds enables some differentiation of the flow into different vessel structures, such as capillaries, venules and arterioles, although one cannot expect a one-to-one correlation between flow speed and vessel type. By separating perfusion into speed regions, the different vascular effects that result from thermal stimulation can be studied separately. Therefore, our results provide an opportunity for deeper insight into endothelial and neurovascular function and for a better understanding of how microcirculation responds to different levels of thermal stimulation in human skin microcirculation.

On the other hand, oxygen saturation represents a dynamic balance between the O_2_ supply and O_2_ consumption in the capillary, arteriolar, and venular beds. It is generally believed that local oxygen saturation is closely related to blood flow perfusion. Previous studies have shown an exponential correlation between oxygen saturation and blood flow. When the flow speed increases, the capillary transit time decreases, resulting in an instantaneously lowered oxygen extraction and an increased overall blood saturation level[15]. However, because blood flow components can be divided into different speed-resolved components, there is still no evidence of the relationship between oxygen saturation and speed-resolved blood perfusion.

This data descriptor outlines a dataset of speed-resolved blood perfusion and oxygen saturation signals recorded locally at the stimulating point before and after stimulation with different levels of thermal stimulation, as well as classical LDF data of the control points and RR interval data of the subjects. It was compiled for our related work[16] with parts of the dataset used in study on complexity changes of local speed-resolved blood perfusion and the relationship between local speed-resolved blood perfusion and the heart rate. The results[16] indicated that different thermal stimulation resulted in different changes in the complexity area index of speed-resolved blood perfusion. Overall, this dataset can be used to study the relationship between local oxygen saturation and speed-resolved blood flux in response to different levels of thermal stimulation. In addition, our present dataset is useful as a reference dataset for various studies on normal or abnormal blood perfusion signal patterns that are common. Because both DRS and LDF have limited penetration depth, the measurements shown here will be restricted to only the superficial regions of the skin.

## Methods

These methods are expanded versions of descriptions in our related work[16, 17].

### Participants and design

Sixty healthy subjects aged 18 to 60 years were included in this study. Please refer to Data 4 for more details[18]. None of the participants reported neurological, psychiatric, or other brain-related diseases. None of the subjects were taking any medication that would affect cardiovascular or autonomic regulation. Alcohol, tea and coffee intake were prohibited for at least 24 hours prior to measurements. This study was conducted according to the Helsinki declaration, all participants were informed of the experimental procedure, and all patients provided written consent prior to the experiment. Written consent was approved by the Institutional Research Ethics Boards of Acupuncture & Moxibustion of the China Academy of Chinese Medical Sciences.

All recruited participants completed the measurements between 9 a.m. and 5 p.m. Subjects in the thermal stimulation group received a total of 4 different levels of stimulation corresponding to 38°C, 40°C, 42°C and 44°C. The order of thermal stimulation was randomly determined for each patient. The experimental protocol was as follows: All experiments were carried out in a quiet, temperature-controlled (24-26°C) laboratory. After a period of cardiovascular stability (30 min), a baseline recording was made for 30 min. Then, the test subjects were stimulated using a thermal stimulation EPOS (Perimed AB, Stockholm, Sweden) for 30 min followed by a 30-min rest period. For the blank control group (BC group), the measurement process was the same as that for the temperature stimulation group (TS group), except that the probe was not heated to keep the subjects in a resting state so that the blood flow parameters of the subjects could be obtained in a resting state. The thermal stimulation point was located on the anterior aspect of the forearm, between the tendons of the palmaris longus and the flexor carpi radials, 5 B-cun proximal to the palmar wrist crease. Control point 1 was located 2 B-cun proximal to the palmar wrist crease, and control point 2 was located 8 B-cun proximal to the palmar wrist crease[19]. The recording points are shown in Figure 1.

**Figure 1.**
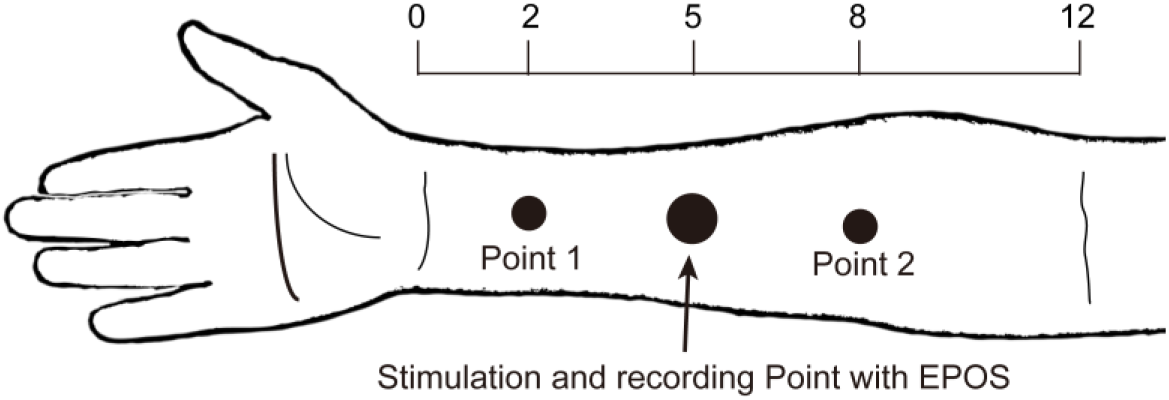
The thermal stimulation point and control point in the right forearm. The thermal stimulation point was located on the anterior aspect of the forearm, between the tendons of the palmaris longus and the flexor carpi radials, 5 B-cun proximal to the palmar wrist crease. Control point 1 was located 2 B-cun proximal to the palmar wrist crease, and control point 2 was located 8 B-cun proximal to the palmar wrist crease. The figure is modified from our related work.

### Protocol for the blood perfusion and oxygen saturation measurements

During the recording session, subjects were placed in a supine position, and their forearms were fixed with a vacuum pillow (AB Germa, Kristianstad, Sweden). Microcirculation was measured by a PeriFlux 6000 Enhanced Perfusion and Oxygen Saturation (EPOS; Perimed AB, Stockholm, Sweden) system with a 3-Hz sampling rate according to the previously described recording protocol[8, 20]. All of the measurement processes were set up in the EPOS management system. During measurements, the EPOS flat probe was fixated using double-sided adhesive tape (PF 105-1, Perimed AB, Stockholm, Sweden). The EPOS system was used for a multimodal assessment of the microcirculation using diffuse reflectance spectroscopy (DRS) and LDF. Using the EPOS system, data for both speed-resolved blood perfusion and oxygen saturation signals were obtained simultaneously. The blood perfusion of the stimulated point was evaluated for three speeds (V1, <1 mm/s; V2, 1-10 mm/s; and V3, >10 mm/s). The probe combined both a laser Doppler probe and a thermostatic probe at the stimulated point, which allowed for controlled, consistent heating of the skin area under the probe surface.

The control points are also located on the midline of forearm, which is collinear with the stimulus point and equidistant from the stimulus point. The classic LDFs of both control points (points 1 and 2) were recorded by a PeriFlux 5000 (Perimed AB, Stockholm, Sweden) system with a 64-Hz sampling rate. The method used to record the blood perfusion flux signal is described in our previous studies[21–23]. Thus in the control points, just conventional blood perfusion was recorded and without any stimulation

Both the thermal stimulation point and control points are shown in Figure 1. The temperature recordings, oxygen saturation recordings, speed-resolved perfusion and classic LDF of the control points for one case subject are shown in Figure 2. For the speed-resolved perfusion results recorded by PF6000, the data were exported directly from the EPOS system in MATLAB format; for the control point results, which were recorded by PF5000, the data were opened with PeriSoft for Windows (version 2.5.5, Perimed, Sweden) and then exported in the txt format. Finally, the data were saved in MATLAB (2015b, MathWorks, Natick, Massachusetts, USA) and Excel format.

**Figure 2.**
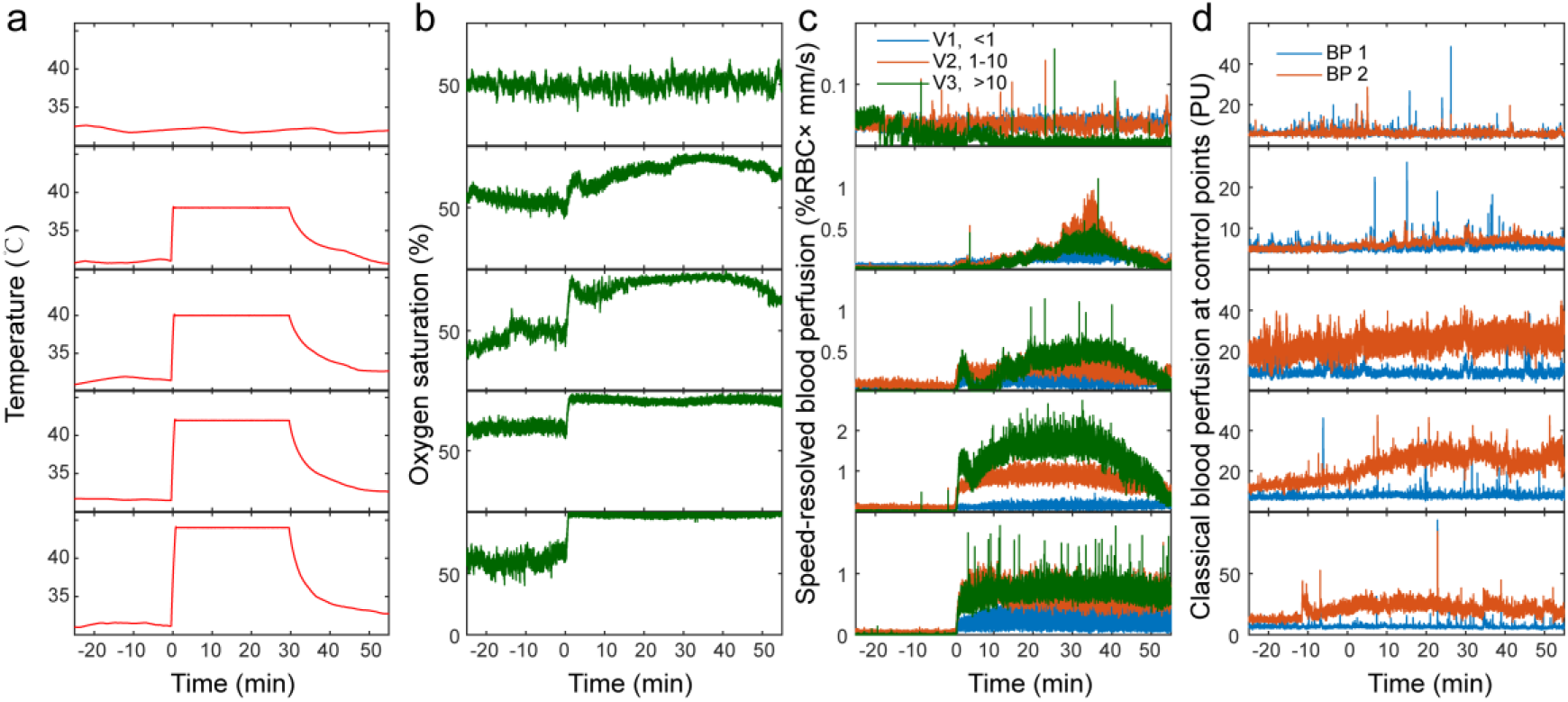
Example of raw dataset. (a) The temperature under different temperature. (b) The oxygen saturation under different temperature. (c) The speed-resolved blood perfusion at the stimulation and recording point under different temperature. (d) The classical blood perfusion at control points under different temperature. V1, low speed, <1 %RBC × mm/s; V2, mid-speed, 1-10 %RBC × mm/s; and V3, high speed, >10 %RBC × mm/s. BP1, blank control point 1; BP2, blank control point 2.

### Electrocardiogram measurement protocol

The ECG was measured by the NeurOne system (NeurOne, MEGA electronics Ltd, Finland). Subjects were directed to lie down on a comfortable bed, and 3 adhesive Ag/AgCl electrodes (3M, Shanghai, China) were applied to the chest for ECG recording at a sampling rate of 1000 Hz[21, 24–26]. The raw data were saved as Data 5 in EDF format, the files were imported into Kubio HRV software (Kubios HRV Premium 3.0.0, Kubios oy, Kuopio, Finland) in EDF format, and then, the 50-min RR interval time series were extracted and saved in mat format.

### Data records

The dataset can be found on the figshare[18]. Data 1, Data 2 and Data 3 folders contain both matlab and excel format files each. Data 5 contains ECG raw data in EDF format (the data structure is shown in Figure 3). The names of the files under each folder are shown in Table 1, Table 2 and Table 3. The mat file in the Data 1 folder contains 4 matrix elements: oxygen, perfusion, temperature and time. The matrix perfusion element is made up of three columns: v1(<1 %RBC × mm/s), v2 (1-10 %RBC × mm/s) and v3(>10 %RBC × mm/s). The oxygen, perfusion and temperature elements are synchronized in time with a 3-Hz sampling rate and are shown in Figure 2a, b, and c. The mat file in the Data 2 folder contains 2 matrix elements, namely, BP_1 and BP_2, from classical LDF recorded at control points with a 64-Hz sampling rate, and these data are shown in Figure 3d. The mat file in the Data 3 folder contains 2 matrix elements: RRs and T_RRs. The RRs represent the R-R time series of the ECG, and the T_RRs represent the time index. The samples were extracted from ECG recordings with a NeurOne system with a 1000-Hz sampling rate. Data 4, comprising data pertaining to the subjects’ characteristics, is presented in CSV format.

**Table 1.** The file names of Data 1 (recorded by EPOS). TS, thermal stimulation; BC, blank control.

| TS dataset name |  |  |  |  | BC dataset name |  |
| --- | --- | --- | --- | --- | --- | --- |
| ID | 38℃ | 40℃ | 42℃ | 44℃ | ID | Dataset |
| T_070 | T6_070_38.mat | T6_070_40.mat | T6_070_42.mat | T6_070_44.mat | T_100 | T6_100.mat |
| T_071 | T6_071_38.mat | T6_071_40.mat | T6_071_42.mat | T6_071_44.mat | T_101 | T6_101.mat |
| T_072 | T6_072_38.mat | T6_072_40.mat | T6_072_42.mat | T6_072_44.mat | T_102 | T6_102.mat |
| T_073 | T6_073_38.mat | T6_073_40.mat | T6_073_42.mat | T6_073_44.mat | T_103 | T6_103.mat |
| T_074 | T6_074_38.mat | T6_074_40.mat | T6_074_42.mat | T6_074_44.mat | T_104 | T6_104.mat |
| T_075** | T6_075_38.mat | T6_075_40.mat | T6_075_42.mat | T6_075_44.mat | T_105 | T6_105.mat |
| T_076 | T6_076_38.mat | T6_076_40.mat | T6_076_42.mat | T6_076_44.mat | T_106 | T6_106.mat |
| T_077 | T6_077_38.mat | T6_077_40.mat | T6_077_42.mat | T6_077_44.mat | T_107 | T6_107.mat |
| T_078 | T6_078_38.mat | T6_078_40.mat | T6_078_42.mat | T6_078_44.mat | T_108 | T6_108.mat |
| T_079 | T6_079_38.mat | T6_079_40.mat | T6_079_42.mat | T6_079_44.mat | T_109 | T6_109.mat |
| T_080 | T6_080_38.mat | T6_080_40.mat | T6_080_42.mat | T6_080_44.mat | T_110 | T6_110.mat |
| T_081 | T6_081_38.mat | T6_081_40.mat | T6_081_42.mat | T6_081_44.mat | T_111 | T6_111.mat |
| T_082 | T6_082_38.mat | T6_082_40.mat | T6_082_42.mat | T6_082_44.mat | T_112 | T6_112.mat |
| T_083 | T6_083_38.mat | T6_083_40.mat | T6_083_42.mat | T6_083_44.mat | T_113 | T6_113.mat |
| T_084 | T6_084_38.mat | T6_084_40.mat | T6_084_42.mat | T6_084_44.mat | T_114 | T6_114.mat |
| T_085 | T6_085_38.mat | T6_085_40.mat | T6_085_42.mat | T6_085_44.mat | T_115 | T6_115.mat |
| T_086 | T6_086_38.mat | T6_086_40.mat | T6_086_42.mat | T6_086_44.mat | T_116 | T6_116.mat |
| T_087 | T6_087_38.mat | T6_087_40.mat | T6_087_42.mat | T6_087_44.mat | T_117 | T6_117.mat |
| T_088 | T6_088_38.mat | T6_088_40.mat | T6_088_42.mat | T6_088_44.mat | T_118 | T6_118.mat |
| T_089 | T6_089_38.mat | T6_089_40.mat | T6_089_42.mat | T6_089_44.mat | T_119 | T6_119.mat |
| T_090 | T6_090_38.mat | T6_090_40.mat | T6_090_42.mat | T6_090_44.mat | T_120 | T6_120.mat |
| T_091 | T6_091_38.mat | T6_091_40.mat | T6_091_42.mat | T6_091_44.mat | T_121 | T6_121.mat |
| T_092 | T6_092_38.mat | T6_092_40.mat | T6_092_42.mat | T6_092_44.mat | T_122 | T6_122.mat |
| T_093 | T6_093_38.mat | T6_093_40.mat | T6_093_42.mat | T6_093_44.mat | T_123 | T6_123.mat |
| T_094 | T6_094_38.mat | T6_094_40.mat | T6_094_42.mat | T6_094_44.mat | T_124 | T6_124.mat |
| T_095 | T6_095_38.mat | T6_095_40.mat | T6_095_42.mat | T6_095_44.mat | T_125 | T6_125.mat |
| T_096 | T6_096_38.mat | T6_096_40.mat | T6_096_42.mat | T6_096_44.mat | T_126 | T6_126.mat |
| T_097 | T6_097_38.mat | T6_097_40.mat | T6_097_42.mat | T6_097_44.mat | T_127 | T6_127.mat |
| T_098 | T6_098_38.mat | T6_098_40.mat | T6_098_42.mat | T6_098_44.mat | T_128 | T6_128.mat |
| T_099 | T6_099_38.mat | T6_099_40.mat | T6_099_42.mat | T6_099_44.mat | T_129 | T6_129.mat |

\*\*, abnormal value of oxygen saturation.

**Table 2.**
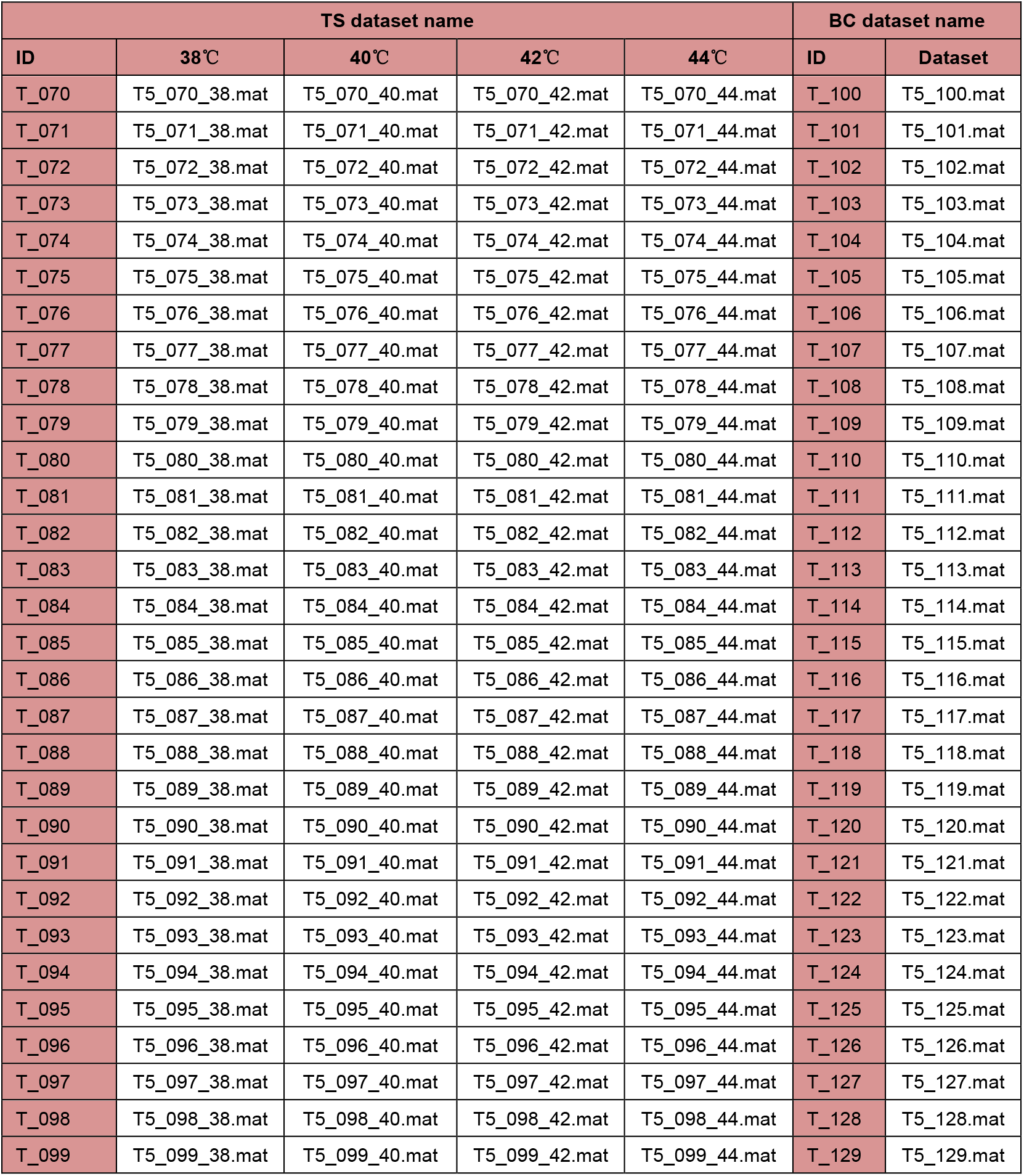
The file names of Data2 (recorded by PF5000). TS, thermal stimulation; BC, blank control

| TS dataset name |  |  |  |  | BC dataset name |  |
| --- | --- | --- | --- | --- | --- | --- |
| ID | 38°C | 40°C | 42°C | 44°C | ID | Dataset |
| T_070 | T5_070_38.mat | T5_070_40.mat | T5_070_42.mat | T5_070_44.mat | T_100 | T5_100.mat |
| T_071 | T5_071_38.mat | T5_071_40.mat | T5_071_42.mat | T5_071_44.mat | T_101 | T5_101.mat |
| T_072 | T5_072_38.mat | T5_072_40.mat | T5_072_42.mat | T5_072_44.mat | T_102 | T5_102.mat |
| T_073 | T5_073_38.mat | T5_073_40.mat | T5_073_42.mat | T5_073_44.mat | T_103 | T5_103.mat |
| T_074 | T5_074_38.mat | T5_074_40.mat | T5_074_42.mat | T5_074_44.mat | T_104 | T5_104.mat |
| T_075 | T5_075_38.mat | T5_075_40.mat | T5_075_42.mat | T5_075_44.mat | T_105 | T5_105.mat |
| T_076 | T5_076_38.mat | T5_076_40.mat | T5_076_42.mat | T5_076_44.mat | T_106 | T5_106.mat |
| T_077 | T5_077_38.mat | T5_077_40.mat | T5_077_42.mat | T5_077_44.mat | T_107 | T5_107.mat |
| T_078 | T5_078_38.mat | T5_078_40.mat | T5_078_42.mat | T5_078_44.mat | T_108 | T5_108.mat |
| T_079 | T5_079_38.mat | T5_079_40.mat | T5_079_42.mat | T5_079_44.mat | T_109 | T5_109.mat |
| T_080 | T5_080_38.mat | T5_080_40.mat | T5_080_42.mat | T5_080_44.mat | T_110 | T5_110.mat |
| T_081 | T5_081_38.mat | T5_081_40.mat | T5_081_42.mat | T5_081_44.mat | T_111 | T5_111.mat |
| T_082 | T5_082_38.mat | T5_082_40.mat | T5_082_42.mat | T5_082_44.mat | T_112 | T5_112.mat |
| T_083 | T5_083_38.mat | T5_083_40.mat | T5_083_42.mat | T5_083_44.mat | T_113 | T5_113.mat |
| T_084 | T5_084_38.mat | T5_084_40.mat | T5_084_42.mat | T5_084_44.mat | T_114 | T5_114.mat |
| T_085 | T5_085_38.mat | T5_085_40.mat | T5_085_42.mat | T5_085_44.mat | T_115 | T5_115.mat |
| T_086 | T5_086_38.mat | T5_086_40.mat | T5_086_42.mat | T5_086_44.mat | T_116 | T5_116.mat |
| T_087 | T5_087_38.mat | T5_087_40.mat | T5_087_42.mat | T5_087_44.mat | T_117 | T5_117.mat |
| T_088 | T5_088_38.mat | T5_088_40.mat | T5_088_42.mat | T5_088_44.mat | T_118 | T5_118.mat |
| T_089 | T5_089_38.mat | T5_089_40.mat | T5_089_42.mat | T5_089_44.mat | T_119 | T5_119.mat |
| T_090 | T5_090_38.mat | T5_090_40.mat | T5_090_42.mat | T5_090_44.mat | T_120 | T5_120.mat |
| T_091 | T5_091_38.mat | T5_091_40.mat | T5_091_42.mat | T5_091_44.mat | T_121 | T5_121.mat |
| T_092 | T5_092_38.mat | T5_092_40.mat | T5_092_42.mat | T5_092_44.mat | T_122 | T5_122.mat |
| T_093 | T5_093_38.mat | T5_093_40.mat | T5_093_42.mat | T5_093_44.mat | T_123 | T5_123.mat |
| T_094 | T5_094_38.mat | T5_094_40.mat | T5_094_42.mat | T5_094_44.mat | T_124 | T5_124.mat |
| T_095 | T5_095_38.mat | T5_095_40.mat | T5_095_42.mat | T5_095_44.mat | T_125 | T5_125.mat |
| T_096 | T5_096_38.mat | T5_096_40.mat | T5_096_42.mat | T5_096_44.mat | T_126 | T5_126.mat |
| T_097 | T5_097_38.mat | T5_097_40.mat | T5_097_42.mat | T5_097_44.mat | T_127 | T5_127.mat |
| T_098 | T5_098_38.mat | T5_098_40.mat | T5_098_42.mat | T5_098_44.mat | T_128 | T5_128.mat |
| T_099 | T5_099_38.mat | T5_099_40.mat | T5_099_42.mat | T5_099_44.mat | T_129 | T5_129.mat |

**Table 3.** The file names of Data 3 (50-min RR interval extracted from Data 5) and Data 5 (ECG signal recorded by NeurOne system). TS, thermal stimulation; BC, blank control.

| TS dataset name |  |  |  |  | BC dataset name |  |
| --- | --- | --- | --- | --- | --- | --- |
| ID | 38°C | 40°C | 42°C | 44°C | ID | Dataset |
| T_070 | T_070_38.mat | T_070_40.mat | T_070_42.mat | T_070_44.mat | T_100 | T_100.mat |
| T_071 | T_071_38.mat | T_071_40.mat | T_071_42.mat | T_071_44.mat | T_101 | T_101.mat |
| T_072 | T_072_38.mat | T_072_40.mat | T_072_42.mat | T_072_44.mat | T_102 | T_102.mat |
| T_073 | T_073_38.mat | T_073_40.mat | T_073_42.mat | T_073_44.mat | T_103 | T_103.mat |
| T_074 | T_074_38.mat | T_074_40.mat | T_074_42.mat | T_074_44.mat | T_104 | T_104.mat |
| T_075 | T_075_38.mat | T_075_40.mat | T_075_42.mat | T_075_44.mat | T_105 | T_105.mat |
| T_076 | T_076_38.mat | T_076_40.mat | T_076_42.mat | T_076_44.mat | T_106 | T_106.mat |
| T_077 | T_077_38.mat | T_077_40.mat | T_077_42.mat | T_077_44.mat | T_107 | T_107.mat |
| T_078 | T_078_38.mat | T_078_40.mat | T_078_42.mat | T_078_44.mat | T_108 | T_108.mat |
| T_079 | T_079_38.mat | T_079_40.mat | T_079_42.mat | T_079_44.mat | T_109 | T_109.mat |
| T_080 | T_080_38.mat | T_080_40.mat | T_080_42.mat | T_080_44.mat | T_110 | T_110.mat |
| T_081 | T_081_38.mat | T_081_40.mat | T_081_42.mat | T_081_44.mat | T_111 | T_111.mat |
| T_082 | T_082_38.mat | T_082_40.mat | T_082_42.mat | T_082_44.mat | T_112 | T_112.mat |
| T_083 | T_083_38.mat | T_083_40.mat | T_083_42.mat | T_083_44.mat | T_113 | T_113.mat |
| T_084 | T_084_38.mat | T_084_40.mat | T_084_42.mat | T_084_44.mat | T_114 | T_114.mat |
| T_085 | T_085_38.mat | T_085_40.mat | T_085_42.mat | T_085_44.mat | T_115 | T_115.mat |
| T_086 | T_086_38.mat | T_086_40.mat | T_086_42.mat | T_086_44.mat | T_116 | T_116.mat |
| T_087 | T_087_38.mat | T_087_40.mat | T_087_42.mat | T_087_44.mat | T_117 | T_117.mat |
| T_088 | T_088_38.mat | T_088_40.mat | T_088_42.mat | T_088_44.mat | T_118 | T_118.mat |
| T_089 | T_089_38.mat | T_089_40.mat | T_089_42.mat | T_089_44.mat | T_119 | T_119.mat |
| T_090 | T_090_38.mat | T_090_40.mat | T_090_42.mat | T_090_44.mat | T_120 | T_120.mat |
| T_091 | T_091_38.mat | T_091_40.mat | T_091_42.mat | T_091_44.mat | T_121 | T_121.mat |
| T_092 | T_092_38.mat | T_092_40.mat | T_092_42.mat | T_092_44.mat | T_122 | T_122.mat |
| T_093 | T_093_38.mat | T_093_40.mat | T_093_42.mat | T_093_44.mat | T_123 | T_123.mat |
| T_094 | T_094_38.mat | T_094_40.mat | T_094_42.mat | T_094_44.mat | T_124 | T_124.mat |
| T_095 | T_095_38.mat | T_095_40.mat | T_095_42.mat | T_095_44.mat | T_125 | T_125.mat |
| T_096 | T_096_38.mat | T_096_40.mat | T_096_42.mat | T_096_44.mat | T_126 | T_126.mat |
| T_097 | T_097_38.mat | T_097_40.mat | T_097_42.mat | T_097_44.mat | T_127 | T_127.mat |
| T_098 | T_098_38.mat | T_098_40.mat | T_098_42.mat | T_098_44.mat | T_128 | T_128.mat |
| T_099 | T_099_38.mat | T_099_40.mat | T_099_42.mat | T_099_44.mat | T_129 | T_129.mat |

**Figure 3.**
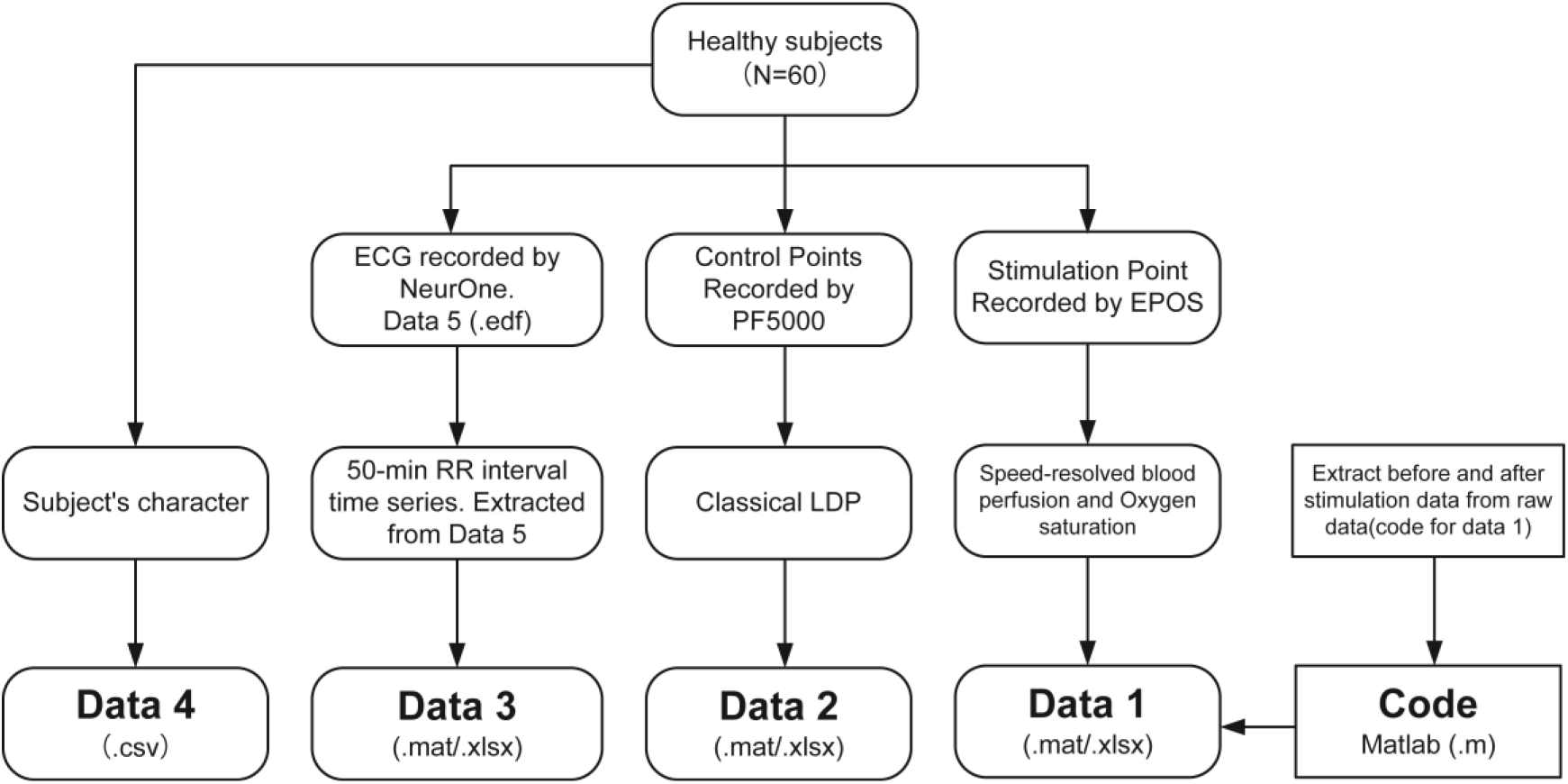
The structure of the database. Data 1, Data 2, Data 3, Data 4 and Data 5 can be found in figshare (https://doi.org/10.6084/m9.figshare.8299343.v4).

### Technical validation

To exclude motion artefacts, the subjects’ forearms were fixed with a vacuum pillow (AB Germa, Kristianstad, Sweden). During the recording, if the haemoglobin signature in the DRS spectra was too weak, the RBC oxygen saturation estimation became unstable. The oxygen saturation estimate was hence discarded for haemoglobin areas that were too small and if the estimated RBC tissue concentration was below 0.1 %[20]. During data preprocessing, any outliers were replaced using the linear interpolation method. By default, an outlier is a value that is more than three scaled median absolute deviations (MADs) away from the median.

In the current dataset, data for oxygen saturation and speed-resolved blood perfusion were obtained simultaneously, and coherence analysis was performed between oxygen saturation and speed-resolved blood perfusion. To calculate the coherence between oxygen saturation and speed-resolved blood perfusion, the coherence value was estimated by the following equation[27]:

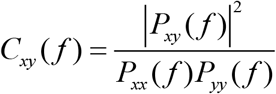

where 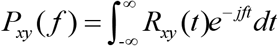, and thus, *P*_*xy*_(*f*) is the Fourier transform of the *R*_*xy*_(*R*_*xy*_ is the cross-correlation of *x* and *y*). This equation obtains the magnitude squared coherence estimate *C*_*xy*_(*f*) of the input signals *x* and *y* using Welch’s averaged, modified periodogram method. The value of *C*_*xy*_(*f*) is between 0 and 1 and indicates how well *x* corresponds to *y* at each frequency. The results (Figure 4) showed that the coherence value between blood flux and oxygen saturation was stable both before and after thermal stimulation.

**Figure 4.**
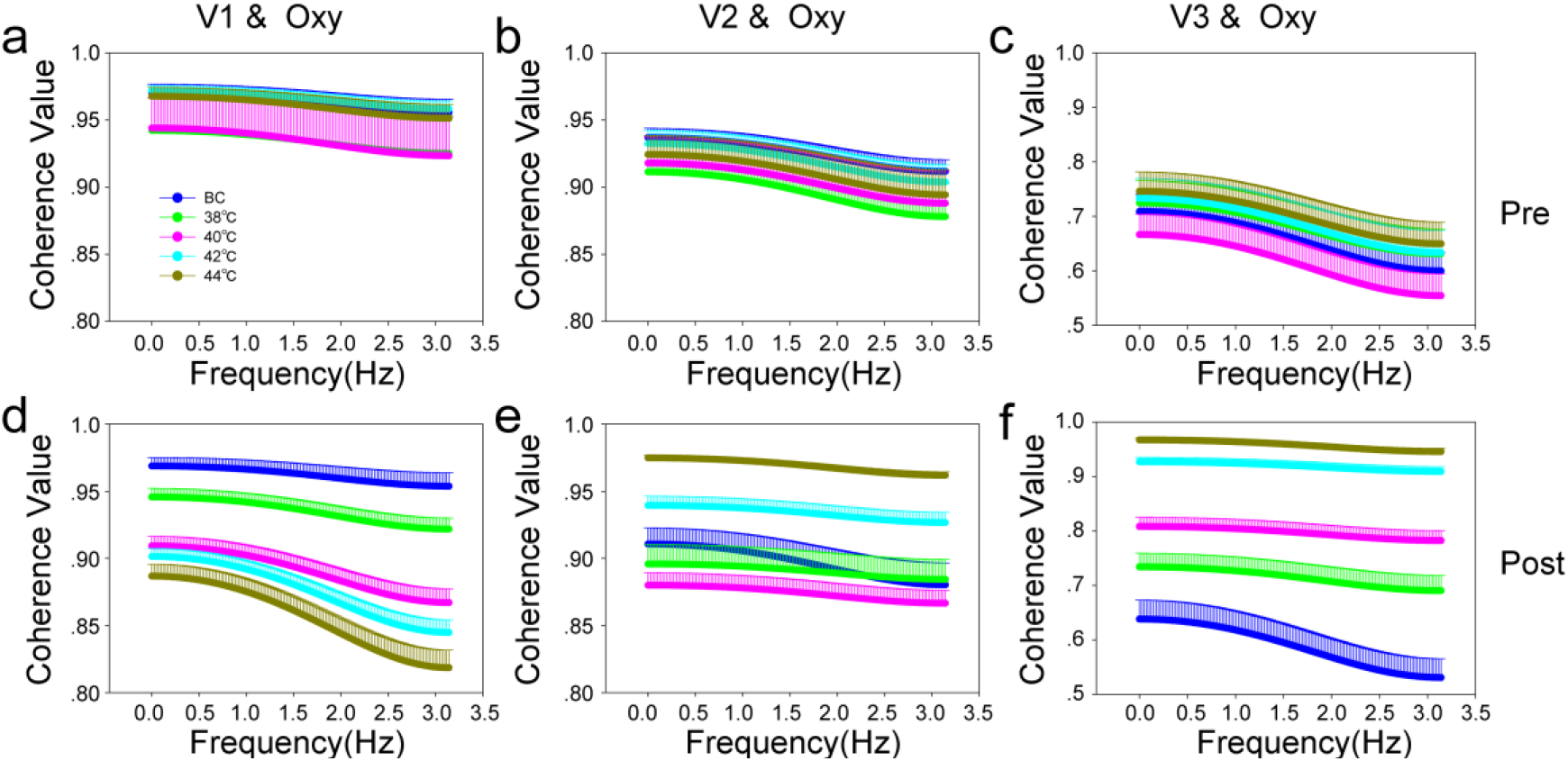
Coherence analysis results between oxygen saturation and speed-resolved blood perfusion. (a) The coherence value between oxygen saturation and V1 blood perfusion before thermal stimulation. (b) The coherence value between oxygen saturation and V2 blood perfusion before thermal stimulation. (c) The coherence value between oxygen saturation and V3 blood perfusion before thermal stimulation. (d) The coherence value between oxygen saturation and V1 blood perfusion after thermal stimulation. (e) The coherence value between oxygen saturation and V2 blood perfusion after thermal stimulation. (f) The coherence value between oxygen saturation and V3 blood perfusion after thermal stimulation. V1, low speed, <1 %RBC × mm/s; V2, mid-speed, 1-10 %RBC × mm/s; and V3, high speed, >10 %RBC × mm/s. Oxy, oxygen saturation. Data are presented as the mean ± sem.

In general, the flow with speeds below 1 mm/s are associated with capillary flow, although some venular flow may also fall into that category. The speeds 1-10 mm/s is foremost constituted by flow in venules, small arterioles and, under some circumstances, veins. The flow speeds above 10 mm/s are primarily related to flow in larger arterioles and other larger vessels situated in the sampling volume. It should be noted, however, that these relationships between flow speed and vessel type may be altered during for example heating provocations or when pressure is applied. Under some conditions, capillaries may act as shunting vessels and then likely contain flow speeds above 1 mm/s, but in that situation those capillaries do not contribute to the nutritive flow. Therefore, when interpreting the data, we should pay attention to the following points: First, the data we get are distinguished by speed, not by blood vessel type; Second, under the condition of thermal stimulation, the standard value of speed to distinguish blood flow may change, no longer 1mm/s and 10 mm/s, and it should be increased appropriately; Third, oxygen saturation are different in different vessel types. It is a potential method to correct speed-resolved blood flow through relationship between oxygen saturation and speed-resolved blood flow.

## Data availability

Guangjun, W., Shuyong, J., Xiaojing, S., Weibo, Z. (2019): A dataset of speed-resolved blood perfusion and oxygen saturation after different thermal stimulation. figshare https://doi.org/10.6084/m9.figshare.8299343.v4. Readers can access codes in our datasets at figshare.com. Among these codes, MATLAB codes named “Code for Data 1.zip” for extract data from raw datasets can be found.

## Acknowledgements

This research was supported by the National Basic Research Program of China (2015CB554502) and the Fundamental Research Funds for the Central Public Welfare Research Institutes (ZZ11-098)

## Author contributions

Wang Guangjun designed and performed the experiment.

Wang Guangjun, Jia Shuyong, Song Xiaojing and Zhang Weibo collected and checked the data.

Wang Guangjun analysed the data.

Wang Guangjun wrote the paper.

All of the authors read, revised and approved the final manuscript

